# A Time Machine for Taxonomy

**DOI:** 10.1101/2024.12.11.627987

**Authors:** Austin Davis-Richardson, Timothy Reynolds

**Affiliations:** One Codex, Inc

## Abstract

The NCBI Taxonomy Database is the primary resource for linking genomic information to taxonomic relationships, widely used across scientific disciplines and critically important to bioinformatics. This database is continuously changing as researchers discover and refine taxonomic relationships. Yet, tracking and comparing past taxonomic states is challenging due to frequent changes and the need to sift through numerous historical snapshots. To address this, we developed the Taxonomy Time Machine: a database for storing many snapshots of a taxonomic tree in a space-efficient manner. We have also created a web-based and programmatic (API) interface to make this data more accessible. This tool is capable of accurately reconstructing taxonomic lineages at any point in the history of the NCBI Taxonomy Database. We demonstrate that this tool is both perfectly accurate and significantly more efficient than loading and querying individual taxonomy snapshots, enabling its use on desktop computers as well as commodity web servers. We have made this tool available on the web (https://taxonomy.onecodex.com) as well as open source under the MIT license (https://github.com/onecodex/taxonomy-time-machine).

## Background & Introduction

The NCBI Taxonomy Database consolidates the taxonomic descriptions of organisms across the tree of life, including viruses, bacteria, archaea, and eukaryotes (Federhen, 2012). These changes to the taxonomic tree are driven by the discovery of new organisms, refinements to taxonomic placement and evolving taxonomic codes used to name taxa and define taxonomic ranks.

Such changes are illustrated by the increasing addition of new taxa derived from metagenomic surveys, the recent renaming of thousands of viral species to conform to Linnean binomial nomenclature (Walker et al., 2021), and the adoption of the Phylum rank by the International Committee on Systematics of Prokaryotes (ICSP^†^; Oren et al., 2021; NCBI, 2021). The availability of genomic data has caused major disruptions especially in the microbial realm, as exemplified by the splitting of *Lactobacillus* into 23 separate genera (Zheng et al., 2020) and the ongoing contributions of the Genome Taxonomy Database (GTDB; Parks et al., 2018). These name changes can apply to long-standing species names as demonstrated by the recently introduced species name *Candidozyma* (née *Candida*) *auris* (Liu et al., 2024).

These continuous updates to the NCBI taxonomy have significant implications, particularly in bioinformatics. Taxonomic identifiers (Tax IDs) do not contain timestamp information meaning that they do not correspond unambiguously to a static lineage, complicating the interpretation of results and potentially confounding meta-analyses. In metagenomics, where taxonomic classification is integral, taxonomic incongruencies can necessitate manual adjustments to compare results between software pipelines (Portik et al., 2022). Approaches that assume a stable 1:1 mapping between Tax IDs and taxonomic lineages can fail, as a single Tax ID in a recent snapshot may correspond to multiple Tax IDs in a previous version. This instability also affects mock community standards, reference lists of pathogens, and biocontrol agents, which may contain outdated names and introduce friction when switching to current valid nomenclature.

While the NCBI offers versioned snapshots of the taxonomy and the NCBI website is capable of redirecting queries for merged or outdated taxa to the current valid taxon, retrieving complete historical lineages or viewing how a taxon has changed over time remains cumbersome. Users must download and parse hundreds of snapshots, which is a time-consuming process. The International Committee on the Taxonomy of Viruses (Lefkowitz et al., 2018)’s historical taxonomy browser (https://ictv.global/taxonomy/history) supports viewing snapshots as far back as 1971 but is limited to viruses. Shen & Ren (2021) developed a tool (TaxonKit) capable of generating a changelog for the NCBI Taxonomy database as well as taxonomic lineages in past versions; it is limited to the command-line.

Here, we introduce Taxonomy Time Machine—an interactive web application, API, and command-line tool designed to enable efficient querying, retrieval, and comparison of taxonomies across time. By storing only incremental changes and employing streamlined queries, our tool reconstructs taxonomic lineages from monthly snapshots across a decade of history beginning when the snapshots first became available in 2014. This approach facilitates historical analyses, improves reproducibility, and provides researchers with a user-friendly interface for examining the temporal dimension of taxonomic data.

## Methods

We retrieved all available snapshots (as of 2024-12-06) ranging from August 2014 to December 2024 (n=123) of the NCBI Taxonomy Database from the NCBI webserver (https://ftp.ncbi.nih.gov/pub/taxonomy/taxdump_archive). The taxonomy snapshots include two tabular flat files: names.dmp (containing rows of Tax IDs, names, and name types) and nodes.dmp (containing rows of Tax IDs, parent IDs, and ranks). By combining the two, one can traverse the tree from a given Tax ID to the root to construct the full taxonomic lineage for a given taxon or conversely iterate down the tree to determine its taxonomic descendants.

Our initial (naive) approach was to store all data from every snapshot of the NCBI taxonomy database by adding a ‘version_date’ column to the table and then resolving queries by first filtering followed by the same tree-traversal process. However, due to the large number of rows in each dump file, storing all rows from every snapshot became impractical. Because of the apparent high redundancy of this data, we developed an approach based on storing only the rows that contained a change since the previous version.

This truncated table, which we call the deduplicated table was generated by iterating over each dump file from the NCBI taxonomy archive in chronological order and only adding nodes that had a change to their ‘name’, ‘rank’, or ‘parent_id’ compared to the same node in the previous version (as matched by tax ID). We added two additional columns to this table in order to store the timestamp of the dump file that the change occurred in as well as the type of change:

• **version_date** - The timestamp of the tax dump file that the change occurred in
• **event_name** - The type of change as defined by the following values
  ∘ **UPDATE** - One or more changes to the parent_id, name, or rank columns
  ∘ **CREATE** - The node did not exist in the previous tax dump file
  ∘ **DELETE** - The node was not found in the current but not the previous tax dump file

By storing only changes, we were able to reduce storage requirements by 98.4% (Figure 1) resulting in a solution that was practical for use on a personal computer or commodity cloud server. A similar approach is used in TaxonKit’s `taxid-changelog` command, however our approach differs in that we do not store the full lineage of each tax ID and the resulting data is stored as an SQLite database which allows for indexing and fast retrieval.

**Figure 1.**
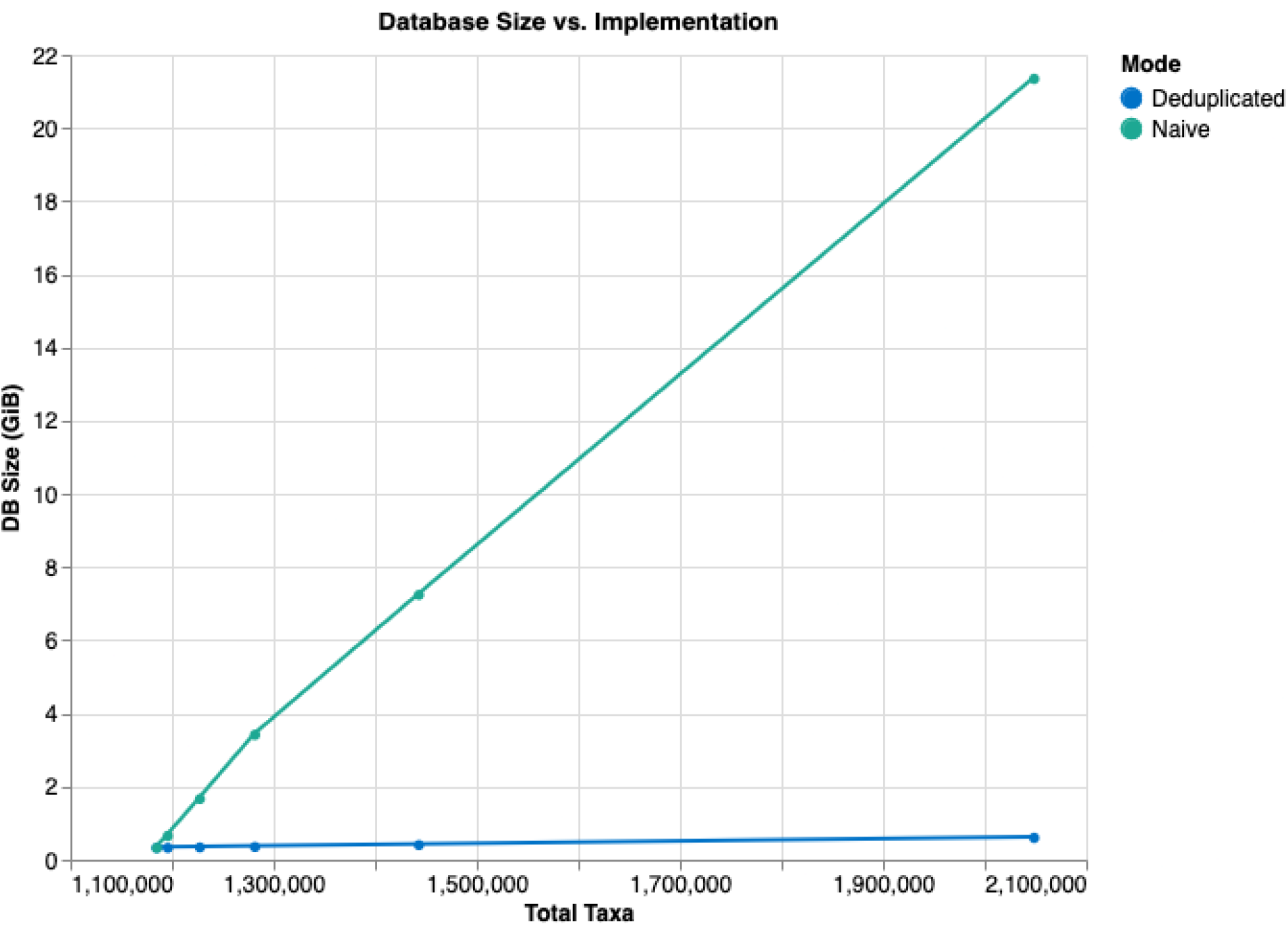
Storage requirements for the taxonomy version database is greatly reduced when redundant (compared to previous version(s)) nodes are skipped as an increasing number of snapshots are added.

We then sought to determine whether it would be possible to query the deduplicated table and generate the same results as the naive approach. We defined the set of queries that we would need to resolve to provide useful information about taxonomies over time:

1. **Query Taxonomic Lineages** (query 1) - Retrieve the taxonomic lineage of a taxon at a given timepoint.
2. **Query Taxonomic Descendants** (query 2) - Retrieve all taxonomic descendants of a taxon (e.g., the species belonging to a genus) at a given timepoint.

Queries are performed against the deduplicated database by traversing “up” the tree from the query taxon to its descendants until racing the root or “down” the tree from the query node to its immediate descendants (child nodes). In order to determine either the lineage of descendants at a given timestamp, rows that were added to the database after that timestamp are excluded from this process. The resultant queries will then return the lineage or descendants as it had been at the timestamp used in the query. An intuitive way to think of this is to imagine filling in deduplicated (removed) rows with their most recent row up until the query date and then traversing the tree as usual.

To assess the accuracy of the deduplicated approach, we exhaustively compared the results of both approaches for every tax ID (n=246,592,264) across every dump file (n=123). The results were compared between both approaches and all resulting lineages and descendants were found to match exactly between the naive and deduplicated approaches.

## Results

We developed an interactive web application to enable searching and browsing taxonomic lineages across snapshots of the NCBI Taxonomy database. The application allows for fuzzy string searching by either name or tax ID. Once a taxon has been selected, the user may view versions of its taxonomic lineage (parents) and children (descendants) across versions as well as see the timestamps for when changes to its taxonomic lineage were introduced. The application also provides an API allowing for programmatic queries.

## Discussion

The temporal dimension of the taxonomic tree cannot be ignored when interpreting scientific names referenced in literature or generated by bioinformatics tools. Our work presents a new tool that allows for fast, user-friendly and computationally efficient access to taxonomic records across multiple timepoints, for the entire tree of life. Future improvements to the tool may include adding earlier snapshots if that data can be provided by NCBI, integrating other taxonomic systems such as the GTDB, and developing automated approaches for normalizing or projecting taxonomies across datasets. We believe this tool will facilitate more accurate and easier interpretation and comparison of taxonomic records. It may be worthwhile to explore applying this process to normalize metagenomic classification results across methods as well as develop automated tools to identify, correct or highlight outdated names used in literature and documentation.

## Availability & Implementation

The application is available at https://taxonomy.onecodex.com, and the source code, including the database used in this manuscript, is available on GitHub (https://github.com/onecodex/taxonomy-time-machine) under the MIT open-source license. The application can be run locally inside a Docker container. The back-end can also be accessed programmatically via a web API or used as a Python module, for example, in a Jupyter Notebook environment.

^†^Formerly known as the International Committee on Systematics of Bacteria

